# Expression of AAACTAC satellite repeats as a long noncoding RNA in the early oocyte of *Drosophila virilis*

**DOI:** 10.64898/2026.08.24.746749

**Authors:** Olivier Vermette, Richy Leonel Mizoy, Jullien M. Flynn

**Affiliations:** Department of Biology, Université Laval, Québec, Canada Institute of Integrative and Systems Biology (IBIS), Québec, Canada

## Abstract

Satellite DNA is long arrays of tandem repetitive DNA located often near the centromeres of chromosomes, whose function, or lack of, has been debated since its discovery. Although situated in heterochromatin, satellite DNA may be expressed as long noncoding RNAs (lncRNAs). Although there are a few examples of satellite lncRNAs being characterized, and functions suggested, how widespread and functionally important they may be for developmental processes is not understood. Here, we take an evolutionary approach to investigate satellite lncRNA expression in *Drosophila spp*. ovaries, a tissue whose development is well-characterized but where satellite expression has only been minimally explored. Using a publicly-available total RNAseq dataset, we find that 118/156 surveyed satellite DNAs were expressed across 10 species, with 33 satellites having high expression over 20 RPM. However, all but two of these expressed satellites (AAACTAC in *D. virilis* and ACAGACAGACAGG in *D. ananassae*) had higher read counts in a sister smallRNA dataset, suggesting that most satellite transcripts primarily serve as precursors for piRNA biogenesis. The two “stand-alone” lncRNAs were highly strand-biased, with 96-97% of the total reads coming from one strand. We further investigated AAACTAC expression with RNA FISH and found the transcript is specifically present in the oocyte nucleus following a dynamic spatiotemporal pattern, with the highest expression in stage 3-5 oocytes. The transcription pattern of AAACTAC is conserved in the three other virilis clade species that contain this satellite DNA. Further, we found expression of unrelated satellites in more distantly related *D. borealis* and *littoralis* both in the oocyte and the nurse cells. Overall, our work identifies a novel lncRNA *AAACUAC* found in the early oocyte nucleus, which is conserved across ∼5 MY of evolution, and is therefore a strong candidate for the discovery of novel functions of satellite lncRNAs in development.

## INTRODUCTION

Satellite DNA is an abundant class of repetitive DNA that repeats in tandem up to millions of times, forming long arrays. Each repeat unit can vary from a few base pairs to hundreds of base pairs, and these sequences mainly cluster in centromeres, pericentromeric regions, or heterochromatin (Shatskikh et al. 2020; Flynn et al. 2020). Although its existence and characteristics are well-established due to its high abundance and ubiquity in eukaryotic genomes, the scope of satellite DNA’s role in the genome is still unclear and debated. Growing evidence shows that satellite DNA can have diverse roles that are biologically meaningful, such as stabilizing the chromatin (Novo et al. 2022), and supporting the chromocenter’s structure (Brändle et al. 2022; Jagannathan et al. 2018). Despite being located in heterochromatin, satellite DNA can be transcribed into noncoding RNAs, especially small regulatory RNAs such as piRNAs (Wei et al. 2021), and long-noncoding RNAs (lncRNAs), which are a broad class of transcripts >200 nt (Shatskikh et al. 2020; Pezer et al. 2011). Like other satellite DNA activities, noncoding satellite DNA transcripts have been proposed to function in a rather circular manner by self-regulating their chromatin state or transcription level (Wei et al. 2021; Biscotti et al. 2015). Satellite-derived lncRNA transcripts have been described to be important for centromere maintenance and chromosome segregation, and the chromatin state of the DNA sequences from which the transcripts originate (Biscotti et al. 2015; Ninomiya et al. 2023). Most studies of satellite lncRNAs have been done in mammals, where transcription is generally permissive and a large portion of the genome is transcribed at a low level, with not all transcripts necessarily having biologically meaningful roles (Palazzo and Koonin 2020; Kapusta and Feschotte 2014). lncRNAs are in emerging study in Drosophila (Shao et al. 2024; Zafar et al. 2023; Poliwal et al. 2026), but to date there have only been a handful of studies on satellite DNA transcripts and with the molecular-level functions not fully understood (Wei et al. 2021; Mills et al. 2019; Kumon et al. 2026; Rošić et al. 2014).

Drosophila is a leading model for studying satellite DNA activities (Flynn and Yamashita 2024; Shatskikh et al. 2020). *D. melanogaster* is the most commonly used for functional studies, but satellite DNAs have been characterized in dozens more species (Wei et al. 2018; de Lima and Ruiz-Ruano 2022; Gebert et al. 2025), and taking an evolutionary approach may reveal novel contexts for satellite DNA activities and functions. The *Drosophila virilis* clade is particularly interesting, having among the largest genomes in the Drosophila genus (Bosco et al. 2007). Approximately 40% of the *D. virilis* genome is composed of three related satellite DNAs (Gall et al. 1971; Gall and Atherton 1974), AAACTAC which is pericentromeric and AAATTAC and AAACTAT which are centromere-proximal (Flynn et al. 2020). AAACTAC is conserved across ∼5 MY (in three other clade species), but the centromere sequences have turned over more rapidly between species (Flynn et al. 2020). Whether the dominant satellite DNA AAACTAC has any functional activity that might permit its abundance and retention in virilis clade genomes has not been explored.

The Drosophila ovary is an excellent tissue to investigate the potential functions of satellite DNA transcripts in development. It is a well-studied developmental system, with the process of oogenesis well-conserved across Drosophila, including in *D. virilis* (Kinderman and King 1973; Calvi et al. 2007). Each ovary contains multiple ovarioles, which are essentially egg assembly lines composed of three cell types: follicle cells, nurse cells, and a single oocyte, whose development unfolds over 14 distinct stages (Bastock and St Johnston 2008). Follicle cells surround the oocyte and nurse cells, forming a membrane that will become the vitelline egg membrane as oogenesis progresses. Nurse cells, numbering 15, are primarily responsible for supporting oocyte development. While follicle cells are transcriptionally active, nurse cells carry out most transcription, producing the majority of products needed throughout all stages of development (Hughes et al. 2018). This includes both coding and non-coding genes transported from nurse cells to the oocyte via the cytoskeleton and ring canals (Becalska and Gavis 2009; Petrella et al. 2007). Notably, the oocyte is generally transcriptionally silent, with chromatin compacted in a karyosome, after meiosis I recombination up until the late stages of oogenesis (Navarro-Costa et al. 2016; Mahowald 1972; King and Burnett 1959). Nurse cells are known to produce piRNA precursors (Wei et al. 2021), as well as retrotransposon transcripts that can be transported to the oocyte (Wang et al. 2018). However, no studies thus far have studied satellite lncRNAs in Drosophila ovaries, nor investigated satellite DNA transcription across a panel of species.

Here, we take an evolutionary approach to characterize satellite DNA-derived lncRNAs expressed in Drosophila ovaries. We start by analyzing a publicly-available total RNAseq dataset of 10 Drosophila species (van Lopik et al. 2023), where we identified two hyper-abundant satellite lncRNAs that do not seem to be processed into piRNAs, one in *D. ananassae* (*CCUGUCUGUCUGU*)_n_ and another in *D. virilis* (*AAACUAC*)_n_. For simplicity, we write the satellite sequences just by their monomer (i.e. *AAACUAC)*. We next used RNA FISH to characterize the expression pattern within the tissue, where the *D. virilis* sequence showed the most interesting pattern of expression in the oocyte nucleus, peaking around stage 5 of oogenesis, which overlaps with the period of complete transcription silencing characterized in *D. melanogaster*. We next perform RNA FISH on other *D. virilis* clade and more distantly-related virilis group species and find some variation in the expression pattern, but that the presence of oocyte nucleus satellite transcripts is conserved in at least five other virilis-group species spanning 11 million years of evolution. Our study provides a benchmark for satellite transcripts across Drosophila (with *D. melanogaster* having among the lowest expression), which highlights the value of studying non-melanogaster species. Finally, our study identifies a very strong lncRNA candidate for future functional studies, which may lead to novel insights on satellite functions and egg development in Drosophila.

## MATERIAL AND METHODS

### Characterization of satellite DNAs in 10 Drosophila species

We initially compiled a comprehensive inventory of satellite DNA sequences across ten *Drosophila* species from different phylogenetic groups, specifically targeting those for which total RNA-seq and small RNA-seq datasets were available: *D. melanogaster*, *D. simulans*, *D. yakuba*, *D. erecta*, *D. ficusphila*, *D. ananassae*, *D. persimilis*, *D. pseudoobscura*, *D. mojavensis*, and *D. virilis*. The inventory was based on satellite DNA discovery using two tools: RepeatExplorer (Novák et al. 2013) which enabled the identification of repetitive sequences, including complex satellite families, while k-Seek (Wei et al. 2014) was employed to detect simple tandem repeats (monomers <=20 bp) and quantify their genomic abundance. We primarily collected satellite sequences from two published articles which have used these approaches: (de Lima and Ruiz-Ruano 2022) for RepeatExplorer and (Wei et al. 2018) for k-Seek. *D. ficusphila* and *D. yakuba* were not included in (Wei et al. 2018), therefore we analyzed published short-read datasets (Bioproject : PRJNA1113679) with k-Seek in order to identify simple satellite sequences to complete these species’ inventory. We applied a log10-normalized abundance threshold greater than 3 to retain only satellite sequences with substantial representation. We excluded mononucleotide repeats from the analysis. In total, we selected 156 satellite sequences to survey across the studied species, corresponding to 11-23 satellite sequences per species (Table S1). We later wanted to study more species in the virilis group to test whether other satellites may be expressed in the ovaries. In the absence of available transcriptomic data for these species, we performed a genomic analysis using whole-genome sequencing (short-read) data from *D. borealis*, *D. littoralis* (Bioproject : PRJNA855919) and *D. montana* (Bioproject PRJNA828433). Satellite sequences were identified and characterized *de novo* using the two satellite DNA identification algorithms. We identified the most abundant satellite DNA in each of these species to test with DNA FISH and then RNA FISH.

## Total RNAseq and small RNAseq data processing

We obtained total RNA-seq (Bioproject : PRJNA937769) and small RNA-seq (Bioproject : PRJNA937774) datasets from Drosophila ovaries generated through large-scale sequencing of multiple Drosophila species by (van Lopik et al. 2023).

## RNA-seq data processing

According to the authors’ protocol, they dissected ovaries from 10–20 flies on ice and extracted total RNA using a TRIzol-based protocol, including chloroform separation and isopropanol precipitation. We processed raw reads to remove adapter sequences (AGATCGGAAGAGC) using Cutadapt v3.2. We applied a filtering step to retain only reads ≥30 nucleotides after adapter removal, ensuring a balance between alignment sensitivity and reliability. Since total RNA-seq libraries were prepared as stranded libraries, we separately quantified reads originating from each strand.

## Small RNA-seq processing

According to the authors’ protocol, they extracted small RNAs from dissociated ovarian cells derived from 75–100 pairs of ovaries using the TraPR Small RNA Isolation Kit (Lexogen). We processed the data with Cutadapt v3.2, including an additional rRNA (TGCTTGGACTACATATGGTTGAGGGTTGTA) depletion step.

## Abundance quantification of satDNA sequences across species

To quantify the relative expression of the different satellite sequences identified across the ten studied species, we generated FASTA files containing the library of tandemly repeated consensus sequences. To allow for mapping of reads that might be overlapping unit boundaries, we included at least three tandem copies of the monomer sequence and ensured that the total array length was at least 50% greater than the read length. Reads were aligned to these reference satellite sequences using Bowtie2 v2.5.3 using the --fr parameter to specify the expected orientation of paired reads (Langmead and Salzberg 2012). We then counted the number of unique reads mapped to each satellite sequence in each biological replicate and each species using Samtools (Li et al. 2009). We then normalized by the total number of reads in each sample to express the relative abundance of satellite transcripts as reads per million (RPM), allowing values to be compared across samples and species.

## Comparison of expression level to control genes

To better contextualize the expression levels of satellite sequences and assess their biological relevance, we included in our analysis a set of reference genes known to be highly expressed in Drosophila ovaries or to be ubiquitously expressed across tissues. These genes were selected from the FlyBase database and comprised early developmental genes involved in embryonic axis establishment *bicoid*, *nanos*, and *gurken* (Kugler and Lasko 2009) a marker gene of germline cell differentiation, bag of marbles (*bam*) (Li et al. 2013), as well as a housekeeping gene associated with protein synthesis, eEF1a1 (Marygold et al. 2017). The coding sequences of these genes were retrieved for each of the ten studied species and incorporated into the alignment pipeline alongside the satellite sequences, thereby generating comparable expression values (in RPM) and providing a reference point to assess the relative abundance of satellite transcripts in Drosophila ovaries.

## Directional orientation of reads

The orientation of reads aligned to each sequence was determined based on SAM flags, following the official Broad Institute documentation (Picard: Explain SAM Flags v3.4.0). Reads carrying, for example, flags 65, 73, 97, and 99 were considered aligned to the forward strand, whereas those carrying flags 81, 83, 89, and 113 were assigned to the reverse strand. For each satellite sequence, the number of reads associated with each orientation was quantified. The relative proportions of forward and reverse transcripts were then calculated and expressed as percentages. This analysis allowed us to assess the presence of a directional bias in the transcription of the sequences under study.

## Fly species and stocks used

We used *D. virilis* 15010-1051.48, *D. americana* ML97.5, *D. novamexicana* 15010-1031.14, *D. lummei* 15010-1011.09, *D. littoralis* 15010-1001.11, *D. borealis* 15010-0961.00, *D. montana* van-08, and *D. melanogaster* Canton-S. Flies were maintained at room temperature or 22°C on standard cornmeal-based media.

## Tissue fixation and probe hybridization

We dissected the ovaries from Drosophila females and fixed them in 1 ml of a solution containing 4% EM-grade formaldehyde (FA) in 1x nuclease-free PBS for 30 minutes at room temperature (RT). After fixation, the ovaries were washed twice with 1 mL of RNase-free 1x PBS for 5 minutes each. They were then permeabilized overnight at 4 °c using 1 mL of RNase-free 70% ethanol. To remove the ethanol, we washed the samples with 1 ml of Wash buffer, which consists of 2X SSC, 10% deionized formamide. The DNA probes for hybridization were synthesized by Integrated DNA Technologies and were labelled at the 5’ end with either Cy3 or Cy5 (see Table S2 for the full list of probe sequences).

We then added 1 µL of the 10 µM probe solution to 99 µL of hybridization solution (2X saline-sodium citrate (SSC), nuclease-free, final concentration; 100mg/mL dextran sulfate (Sigma, D8906), final concentration; 1 mg/ml yeast tRNA (Sigma, R8759), final concentration; 2 mm vanadyl ribonucleoside complex (NEB, Catalogue Number S142), final concentration; 0.5% RNAse-free BSA (Ambion, AM2618), final concentration; 1 ml deionized formamide; and nuclease-free water to a final volume of 10 ml). We adopted multiplex probe hybridization to reduce nonspecific hybridization of similar probe sequences (e.g. AAACTAT and AAACTAC). The microtubes containing the hybridization solution were then sealed with Parafilm on the cap, placed in a 37°C water bath, and protected from light for at least 18 hours (or overnight). The ovaries were then washed with 1 mL of Wash buffer three times, each for 20 minutes, for a total of 1 hour, in the 37°C water bath. After removing all wash buffer from the last wash, we added 40 µL of ProLong™ Gold mounting solution with DAPI to the microtubes containing the ovaries, and then left the sample to absorb the DAPI for at least 2 hours at 4°C in the dark.

## Mounting the ovaries on slides and observing the tissues

After resting in the mounting media at 4°C, the ovaries were placed between a glass slide and a coverslip using disinfected forceps, sealed with nail polish, and stored again at 4°C in the dark. The confocal microscope used was the Leica SP8 with LAS X software. We mainly used the hybrid (HYB) mode with the three channels (DAPI, Cy3, Cy5): DAPI in PMT mode and the others in HYB mode. We captured at least 10 images per ovary across different oocyte maturation stages, with or without specific signals.

We also used the wide-field microscope Revolve by Echo to capture images of ovaries for qualitative assessment for the following samples: the forward strand of the satellite DNAs found in *D. littoralis, D. lummei, D. montana* and *D. borealis*, and the reverse strand of the lncRNA found in *D. ananassae*.

## RNase A treatment

We conducted an RNase treatment prior to probe hybridization to verify that our RNA-FISH signal is not from non-specific hybridization with DNA. The RNase solution contained 5 mg/mL RNase A, 1 M Tris, 3 M NaCl, and 0.5 M EDTA, with all components at their final concentrations. This solution was applied to the ovaries at 37°C just before hybridization and incubated for 2 hours. Afterwards, the RNase was removed with three 10-minute washes with Washbuffer at 37°C.

## Expression score quantification

We analyzed the images in Fiji, in which the different channels of interest were separated, and then the oocyte signal was categorized into four levels: none, weak, medium, and high (Figure 2A). Then, the data files were opened in RStudio, and each category was then counted for every oocyte development stage. Furthermore, we calculate the expression score by summing the product of each stage’s intensity and its associated score (0 = none, 1 = weak, 2 = medium, 3 = high).

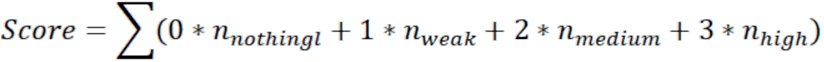

**Figure 1.**
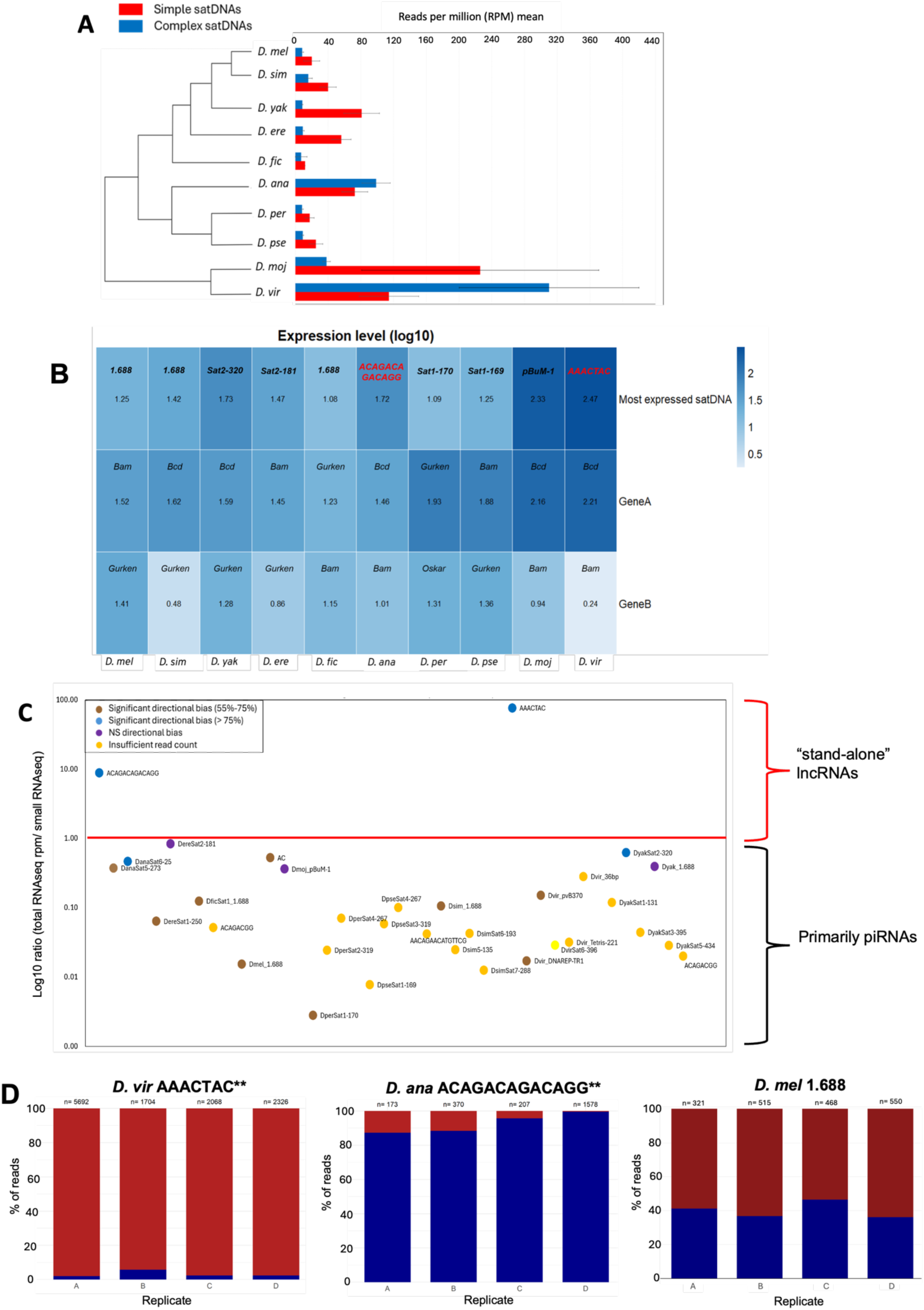
Two stand-alone lncRNAs are highly expressed in a strand-biased manner in the ovaries of two Drosophila species: AAACTAC (*D. virilis*) and ACAGACAGACAGG (*D. ananassae*). A) Total ovarian expression of satellite DNAs in each of 10 Drosophila species, segregated by simple satellites (unit length <= 20 nt) and complex satellites (unit length > 20 nt). B) Comparison of the most highly expressed satellite DNA in each species (top row) to a similarly highly expressed well-known ovarian development gene (middle row) and a lower expressed development gene (bottom row). C) Comparison of total vs. small RNA seq expression level and strand representation of various satellite DNAs expressed across the 10 species. The two stand-alone lncRNAs have much higher expression in the total RNAseq and a strong strand bias. D) Strand bias of the two stand-alone lncRNAs: red represents the forward strand (equal to the sequence written above) and blue represents the reverse strand (reverse complement to the sequence written above). 1.688 (*D. melanogaster*), which is expressed primarily as piRNAs, is shown as a comparison.

**Figure 2.**
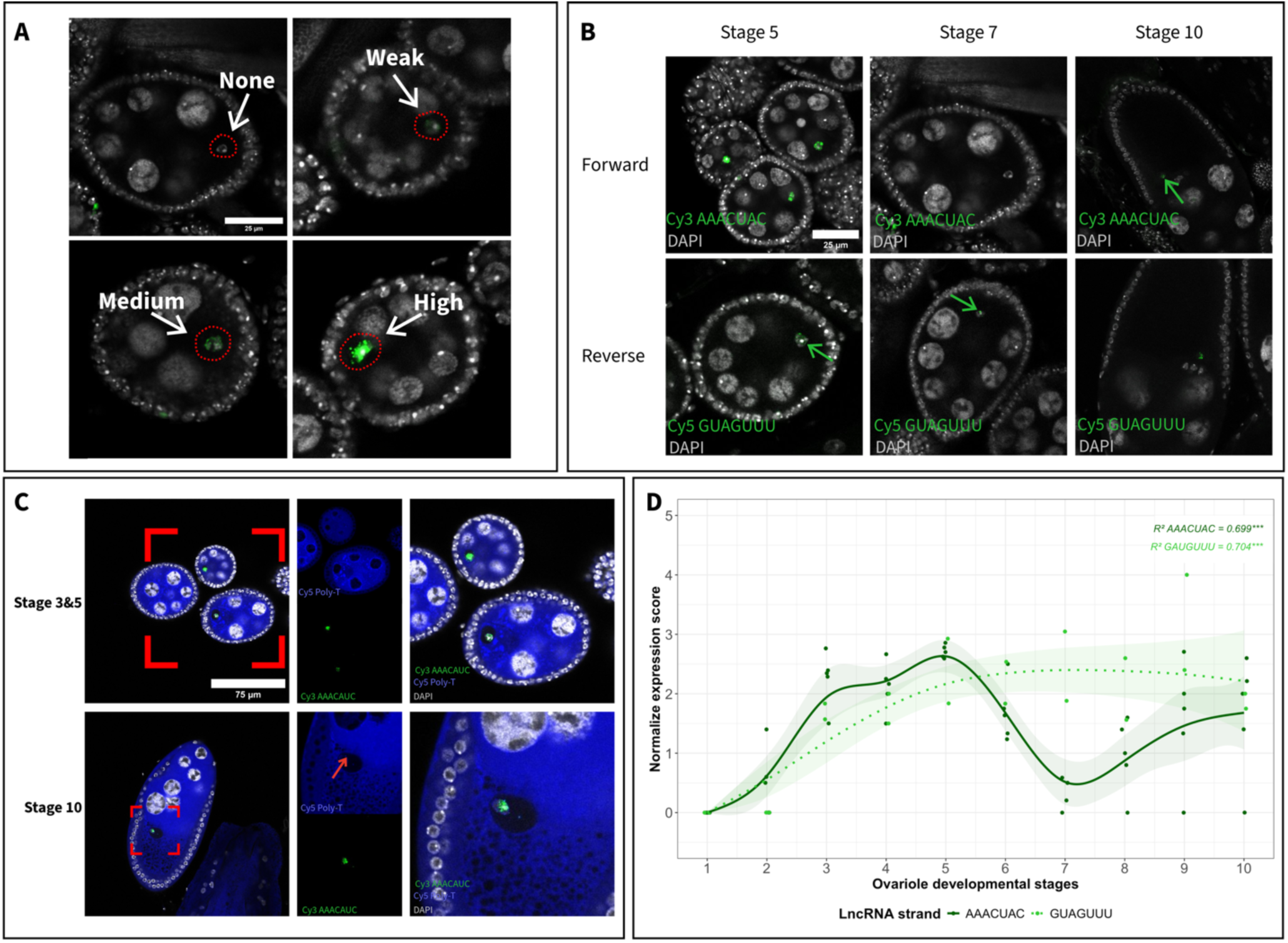
Expression pattern of the *AAACUAC* lncRNA in *D. virilis* ovaries. A) The expression score categorization (non, weak, medium, high) reference for the oocyte was utilized universally for all transcripts and across all species, encompassing all developmental stages. Notably, the majority of ovaries in this reference are in their early stages, as the transcription of most transcripts predominantly occurs during this period. B) b’) *AAACUAC* expression in the oocyte for stages 5-7-10. b’’) *GUAGUUU* (reverse strand) expression in the oocyte for stages 5-7-10. C) *AAACUAC* expression in the oocyte for stages 5-7-10 multiplexed with Poly-A tail probe (T)*_30_*. D) GAM model quantification of the normalized expression score of *AAACUAC* forward (solid green line) and reverse (dotted green line) in *D. virilis*. The total number of observed ovarioles is 1018 for both strands (43 ovaries). Forward strand has a total of 528 (27 ovaries): Stage 2 = 8, Stage 3 = 76, Stage 4 = 18, Stage 5 = 138, Stage 6 = 48, Stage 7 = 110, Stage 8 = 66, Stage 9 = 38, Stage 10 = 26. The points represent the average of the four slides used. Reverse strand has a total of 490 (16 ovaries): Stage 2 = 2, Stage 3 = 48, Stage 4 = 10, Stage 5 = 136, Stage 6 = 57, Stage 7 = 107, Stage 8 = 76, Stage 9 = 28, Stage 10 = 26. The points represent the average of the four slides used.

Subsequently, a normalization of the scores was performed by dividing each stage’s score by the number of occurrences of that stage, with an additional factor of +1 to avoid division by zero.

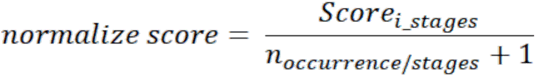

## Colocalization analysis

The 3D colocalization tests were performed on Fiji with the Jacop2 plugin (Aditya et al. 2025; Bolte and Cordelières 2006). Where we calculated the Pearson correlation factor (Bolte and Cordelières 2006; Adler and Parmryd 2010), the MOC, Mander’s coefficients (M1 and M2) (Adler and Parmryd 2010), the ICQ the coefficient (Bolte and Cordelières 2006; Dunn et al. 2011), Li plots, cytofluorograms (Bolte and Cordelières 2006), and the Van Steensel plot were computed. Since all scores, taken individually, are biased to some extent, we needed to calculate and consider all these scores, which allows us to draw a more confident conclusion about the 3D spatial colocalization of the transcripts.

We have also built an analysis macro in Fiji to automate all the colocalization tests for each ovarian stage (https://github.com/OliV-cell/Colocalisation_analysis). Then, the colocalization data were imported and analyzed in R 4.5.2 and Julia 1.12.5 (Aditya et al. 2025; Adler and Parmryd 2013; Miura and Sladoje 2019). Additionally, we have measured a threshold using Rényi’s entropy (Sahoo et al. 1997) to isolate oocyte expression from background noise during the colocalization analysis. Additionally, Coste’s randomization was not performed to limit the analysis’ computational cost, and we conclude that this step was redundant given our data type and the thresholding algorithm.

## Statistical analysis

The statistical analyses were conducted in RStudio version (2026.07.1-147.exe). The models we used were: a linear regression model (LM) for the 3D ICQ sense (4-spline), a GLM (ord-bêta distribution) for M1 3D forward transcript score (5-spline) and a GLM Gamma for the M2 3D forward transcript score (5-spline). Additionally, we fitted GAM (Generalized Additive Model) models with a Gaulss distribution, a type of Gaussian distribution used to calculate the variance of the mean as part of the modeling process, to model the evolution of the expression score in relation to the developmental stages of the oocyte. This type of model is commonly used with non-linear data responses, and for us, it was the case; our expression score in relation to our developmental stages is indeed non-linear. All detailed models, plugins, and packages used are provided in the supplementary information. An ANOVA comparing the different fits for each species was conducted, along with the post hoc Tukey test, to assess whether the GAM models, for each satellite transcript and for each species, were statistically different from each other and for each specific stage. The normality of each model was assumed since they each follow the previously mentioned Gaulss distribution.

## RESULTS

### Evolutionary survey reveals that satellite DNAs are broadly and variably expressed in Drosophila ovaries

First, we analyzed a public dataset containing total RNAseq of ovaries for 10 Drosophila species (van Lopik et al. 2023) (Figure 1A). We quantified the expression of a total of 156 different satellite DNAs, and were able to detect expression for 118 of them, with all species having at least one satellite sequence expressed at 10 RPM or more (Figure 1A, Table S3). 33 satellite sequences were expressed very highly, at 20 RPM or more, and *D. melanogaster* and *D. ficusphila* were the only two species that did not have any individual satellites expressed above this threshold. Notably, the well-characterized 1.688 satellite sequence was the highest expressed in *D. melanogaster*, followed by AAGAG and Rsp. 1.688 was also expressed in *D. simulans, D. yakuba, D. erecta, D. ficusphila,* and *D. ananassae*, demonstrating the broad conservation of this sequence’s expression. The most highly expressed satellite in each species corresponded to the satellite with the documented highest or second highest genomic abundance in 8/10 cases (Table S4). The two exceptions were 1.688 in *D. melanogaster* and *D. simulans*, which was the highest expressed but documented as the 5th or 4th most abundant satellite, respectively.

The representation of simple vs. complex satellite DNA transcripts also varied among the species, with only *D. virilis* and *D. ananassae* having a higher total transcript abundance for simple satellites (unit length <= 20 bp) (Figure 1A). *D. virilis* had the highest level of satellite expression, which was dominated by the abundant pericentromeric simple satellite AAACTAC (Flynn et al. 2020), expressed at 293 RPM on average. *D. mojavensis* had the second highest level of satellite expression, which was dominated by the pBuM satellite, a ∼190 bp previously-described satellite in cactophilic Drosophila (de Lima et al. 2017), expressed at 215 RPM on average.

To add context to our RPM quantifications, we measured the expression of well-known genes expressed in the ovary in the same total RNAseq datasets (Figure 1B, Table S5). Notably, AAACTAC in *D. virilis*, ACAGACAGACAGG in *D. ananassae,* and pBuM in *D. mojavensis* all had higher RPMs than *bicoid* transcripts.

## lncRNAs that are not processed into small RNAs are relatively rare in Drosophila ovaries

Previous studies have found that satellite DNAs can be transcribed as lncRNA precursors before being processed into small RNAs, particularly piRNAs (piwi-interacting RNAs) (Han and Zamore 2014; Wei et al. 2021). In that case, the lncRNAs might just be a transitory molecule that does not have itself an activity. Therefore, we analyzed the associated small RNAseq dataset available from (van Lopik et al. 2023) to quantify the small RNA abundance of each expressed satellite. We then compared the ratio of RPM in total RNAseq vs. small RNAseq (Figure 1C, Table S6). As a conservative interpretation, we considered satellites with a total RNAseq RPM : small RNAseq RPM ratio above 1.0 to be satellites that are expressed as lncRNAs not majorly processed into small RNAs (“stand-alone lncRNAs”). Notably, only two highly-expressed satellite DNAs fit these criteria, AAACTAC in *D. virilis* (ratio 76), and ACAGACAGACAGG in *D. ananassae* (ratio 9). The highly expressed pBuM satellite in *D. mojavensis* was highly expressed as small RNAs (ratio 0.36). We found convincing support that 1.688 is expressed primarily as a small RNA having a ratio of 0.015. Interestingly, the two stand-alone lncRNAs originate from simple satellite DNAs. However, we also found that simple satellite DNAs do form small RNAs in some species, such as AACAGAACATGTTCG in *D. simulans* (ratio 0.042), and ACAGACGG in *D. yakuba* and *D. ficusphila* (ratio 0.02 and 0.051, respectively).

We next quantified the contribution of each of the two possible strands in the total RNAseq dataset to verify if “stand-alone” satellite lncRNAs have a different regime of expression compared to satellite lncRNAs processed into small RNAs. We found that the stand-alone lncRNAs in *D. virilis* and *D. ananassae* were highly strand-biased (96-97% of the reads coming from just one strand), whereas the majority of the lncRNAs processed into small RNAs have both strands represented with less of a bias (Figure 1C, Table S7). The forward strand (the strand containing more As) was expressed predominately for AAACTAC (RNA sequence *AAACUAC*) in *D. virilis*. The reverse strand was expressed predominantly for *D. ananassae* ACAGACAGACAGG (RNA sequence *CCUGUCUGUCUGU*) (Figure 1D). As a comparison, 1.688 in *D. melanogaster* had both strands represented (Figure 1D). Because of our lab’s particular interest in *D. virilis*, we focused on *AAACUAC* expression in *D. virilis* for further analysis, while returning to *D. ananassae* for an evolutionary perspective.

## *AAACUAC* lncRNA is highly expressed in stage 3-5 oocytes, and then again moderately in stage 9-10

We used single-molecule RNA FISH to characterize the expression pattern of the *AAACUAC* in ovaries. Surprisingly, we found that the transcript is most highly expressed in the oocyte nucleus, although there is some expression in occasional follicle cells (Figure 2A). The signal completely disappeared when ovaries were treated with RNase A before the hybridization, indicating a true signal from our RNA (Figure S1). For further analysis, we focused on the expression in the oocyte. We categorically characterized expression at each stage of oogenesis (none, low, medium, high; see methods for details), and found the transcript is highly expressed in the oocyte of stage 3-5 egg chambers, before expression dropping off during stages 6-8 and then returning to moderate levels at stages 9-10 (Figure 2B, D). We could also detect transcripts of the opposite, but less highly expressed strand (*GUAGUUU*), and there was less marked spatio-temporal variability (Figure 2B, D). We also note that from the RNAseq data, the *GUAGUUU* strand only made up 3% of reads, thus we focused most of our analysis on the more abundantly-expressed strand.

The expression of *AAACUAC* in the oocyte nucleus was surprising given the fact that the oocyte of *D. melanogaster* is known to be transcriptionally silent until late oogenesis, and oogenesis is broadly conserved across Drosophila. We therefore used a poly-T probe, which would hybridize to poly-A tails, a universal feature of mRNAs. Over 75% of *D. melanogaster* lncRNAs do not contain poly-A tails (Shao et al. 2024). We found that the poly-A signal was completely excluded from the early oocyte nucleus in *D. virilis* where *AAACUAC* is expressed (Figure 2C). The exclusion pattern of poly-A signal was identical in *D. melanogaster* (Figure S2). This confirms that the lncRNA *AAACUAC* does not contain a poly-A tail, and that poly-A mRNA is not present in the oocyte nucleus during these early stages of development in *D. virilis,* just as in *D. melanogaster*. Nevertheless, some poly-A signal appears in the nucleus of late oogenesis in both *D. virilis* and *D. melanogaster*, which is expected given that the oocyte transcription starts to turn on around stage 9 (Figure S2).

## Related centromere-proximal satellite AAACTAT is expressed moderately in stage 9-10

*D. virilis* has three dominant 7 bp satellite DNAs: AAACTAC (pericentromeric), AAATTAC (centromeric on chromosomes X, 2, 4), AAACTAT (centromeric on chromosomes 3 and 5 and present on the Y and dot chromosome), and one less abundant sequence: AAACAAC, which is present at a single locus (Flynn et al. 2020). We used RNA FISH to verify whether any of these other sequences were expressed. Initial single-plexed experiments showed that the signal for *AAACUAU* and *AAAUUAC* was identical to that of *AAACUAC*. As a control, we did RNA FISH targeting *AAAUUAC* in *D. americana*, which does not contain this satellite DNA in its genome, and we again found hybridization patterns resembling *AAACUAC* (Figure S3). We were therefore concerned that these probes were cross-hybridizing to *AAACUAC*, since there is only one nucleotide difference between them, so we proceeded with multiplex experiments to allow for competition for hybridization between probes, combining the probe targeting *AAACUAC* with a probe targeting either *AAACUAU* or *AAAUUAC*.

In these multiplexed experiments, we could not detect any signal of *AAAUUAC* in *D. virilis* ovaries, thus we concluded it is not expressed. The probe targeting *AAACAAC* cross-hybridized with the poly-A tails of mRNAs (Figure S4), but we could not detect any signal in the oocyte nucleus so we conclude that this sequence is likely not expressed either, at least not in a similar manner as *AAACUAC*. Finally, we found that *AAACUAU* is expressed in the oocyte nucleus, mainly at the later stages of oogenesis (stages 9-10; Figure 3 A,C). 3D colocalization analysis showed that the two satellite transcripts are weakly colocalized and mainly occupy separate nuclear regions. This is reflected in the overlap scores M1 and M2, which measure how much *AAACUAC* volume lies within *AAACUAU*, and vice versa. As shown in Figure 3D-E, the transcripts have a constant but low share of volume: *AAACUAC* comprises about 10% of *AAACUAU*’s volume, while *AAACUAU* makes up roughly 25% of *AAACUAC*’s volume. This indicates that although the transcripts occupy some shared space within the oocyte nucleus, they each have distinct territories. This divergence of territory can also be seen in the decrease in the ICQ score (Figure 3B) as the oocyte matures and the expression of those transcripts enters their second highly expressed step, providing further evidence of the distinct spatial localization of *AAACUAC* and *AAACUAU*.

**Figure 3.**
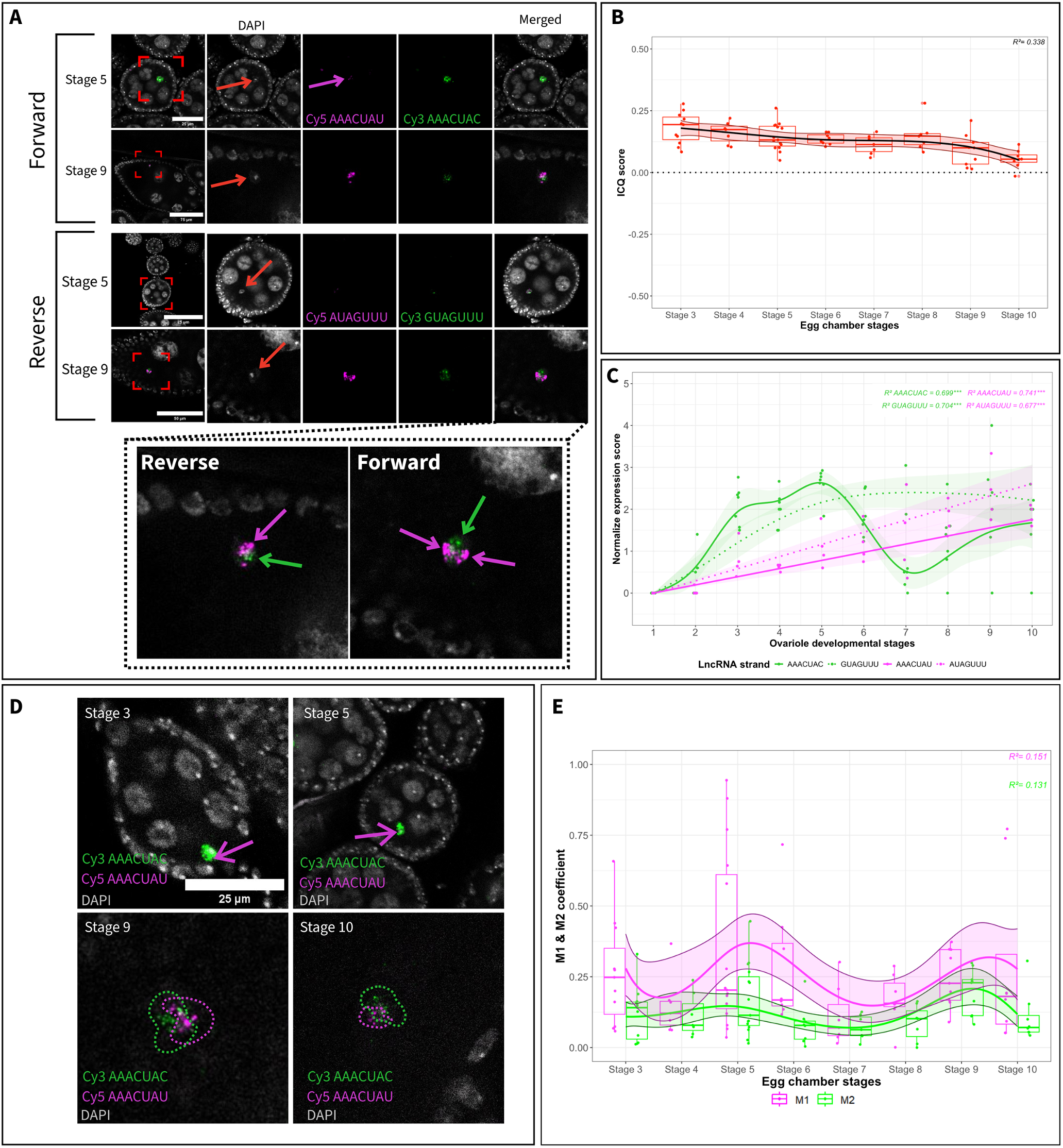
The related centromere sequence AAACTAT is also expressed, mostly in later oocyte development. A) *AAACUAC* and *AAACUAU* expression in the oocyte for stages 5-9 and their reverse-specific expression *GUAGUUU* and *GUAGUUU*) B) linear regression model (LM 4-spline) of the ICQ score through the oocyte development. The toal numbers of observed ovarioles is 72. Stage 3 = 11, stage 4 = 7, stage 5 = 15, stage 6 = 8, stage 7 = 7, stage 8 = 7, stage 9 = 9, stage 10 = 8. C) Quantification of *AAACUAC* forward transcripts (solid green line), *AAACUAU* transcripts (solid magenta line), in the reverse transcripts *GUAGUUU* (dashed green line), and *AUAGUUU* (dashed magenta line). The total number of observed ovarioles is 1018 across both strands (43 ovaries). Forward strand, total of 528 (27 ovaries): Stage 2 = 8, Stage 3 = 76, Stage 4 = 18, Stage 5 = 138, Stage 6 = 48, Stage 7 = 110, Stage 8 = 66, Stage 9 = 38, Stage 10 = 26. The points represent the average of 4 slides used. Reverse strand = 490 (16 ovaries): Stage 2 = 2, Stage 3 = 48, Stage 4 = 10, Stage 5 = 136, Stage 6 = 57, Stage 7 = 107, Stage 8 = 76, Stage 9 = 28, Stage 10 = 26. The points represent the average of 4 slides used. D) *AAACUAC* and *AAACUAU* show forward-specific expression in the oocyte at stages 3, 5, 9, and 10. E) The voxel overlapping score quantifies spatial disparity in the oocyte between the *AAACUAC* and *AAACUAU* forward transcripts. The M1 score uses a Generalized Linear Model (GLM) with an ordinal-beta distribution and a 5-knot spline, while the M2 score employs a Gamma GLM with a 5-knot spline.

## Expression of AAACTAC is conserved in other virilis clade species

The satellite DNA AAACTAC is conserved in the pericentromeric region in the other virilis clade species spanning 4.5 million years, including *D. novamexicana*, *D. americana*, and *D. lummei* (Flynn et al 2020; Figure 5A). We also probed for *AAACUAC* expression in these species and found that expression in the oocyte was largely conserved across all three species (Figure 4B, Table S8). *D. lummei*’s expression pattern was the most different from *D. virilis* and the other species, being expressed significantly more weakly in stages 3-5 and more highly in stage 7, with expression actually peaking at stage 7. *D. novamexicana* showed lower expression than *D. virilis* and *D. americana* at stage 5, and *D. americana* showed higher expression than *D. virilis* at stage 7. Since *D. novamexicana* and *D. americana* possess the satellite AAACAAC centromere-proximally, we probed for its expression as well. We found the same pattern as in *D. virilis*, that the probe cross-hybridized with poly-A tails but was not expressed in the oocyte nucleus with *AAACUAC*.

**Figure 4.**
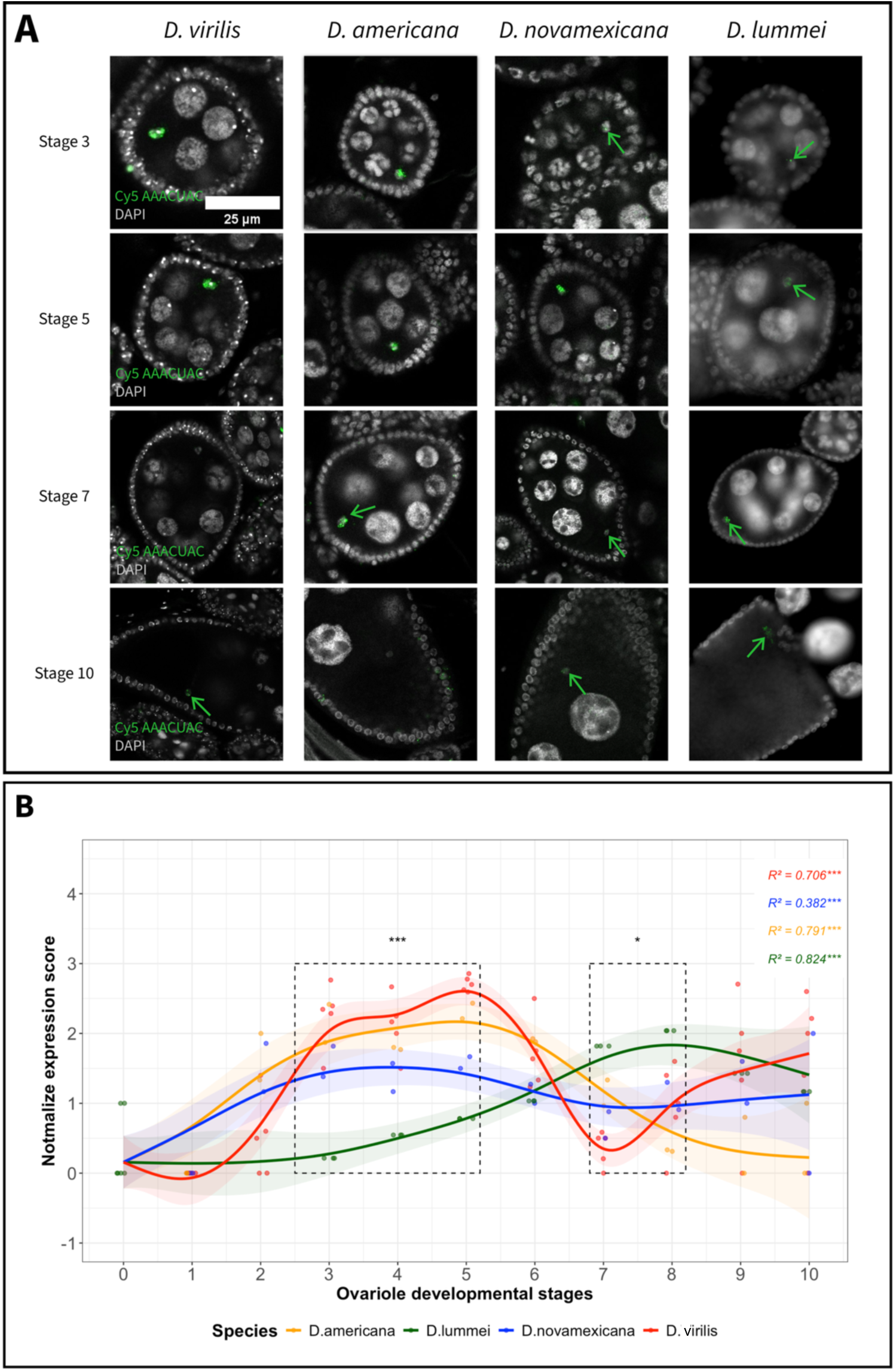
Comparison of AAACTAC expression across the virilis clade A) Expression of *AAACUAC* in *D. virilis*, *D. americana*, *D. novamexicana,* and *D. lummei*, depending on the developmental stage 3-5-7-10. B) GAM model quantification of the normalized expression score for the sense transcript *AAACUAC*, depending on the developmental stage and species, with only *D. lummei* being statistically different. The total number of ovarioles observed for *D. virilis* is 635 (52 ovaries), for *D. americana* it is 285 (20 ovaries), for *D. novamexicana* it is 169 (9 ovaries), and for *D. lummei* it is 288 (20 ovaries). Each point represents the average of each experiment.

**Figure 5.**
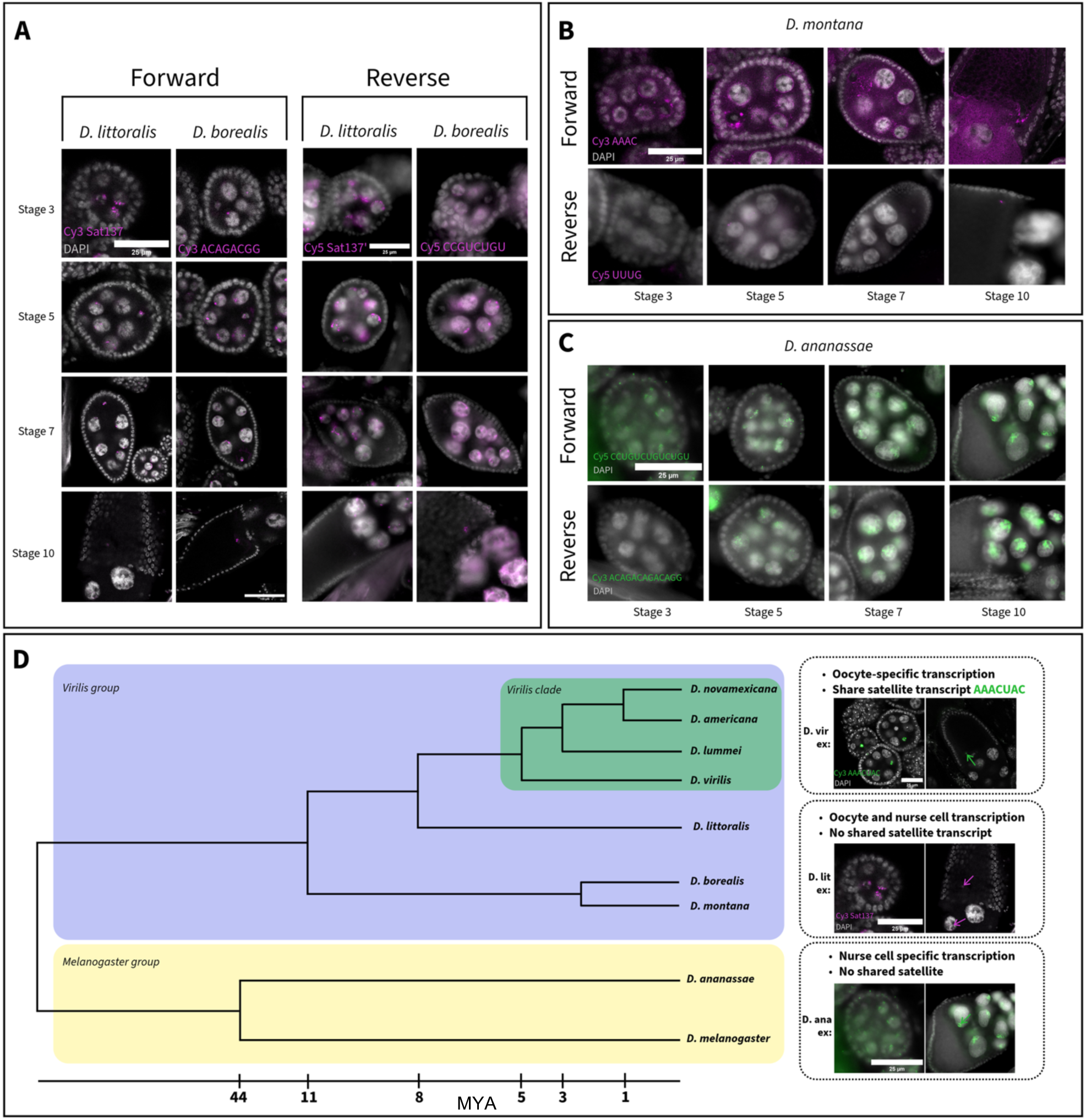
Expression of satellite DNAs outside of the virilis clade A) Hybridization of the complementary probe of the forward transcript of sat137 for *D. littoralis* and ACAGACGG for *D. borealis*, showing the expression within nurse cell and oocyte for all stages. B) Hybridization of the complementary probe of the forward and reverse transcript of AAAC for *D. montana*, the forward probe hybridized with Poly-A tails and we cannot conclude expression. The reverse probe shows weak late-stage expression, specific to the oocyte. Images were taken with the Revolve microscope from Echo. C) Expression of *ACAGACAGACAGG* (forward) and *CCUGUCUGUCUGU* (reverse) in the nurse cells of *D. ananassae* ovaries. D) Phylogenetic tree summarizing the varying expression patterns we found, including 9 Drosophila species (*D. melanogaster* shown for orientation; branch length not to scale).

## Other virilis group species display expression of unrelated satellites in both nurse cells and oocytes

Since satellite lncRNA expression was found to be relatively rare in Drosophila genus ovaries, but conserved in the virilis clade, we decided to expand our RNA FISH experiments to species more distantly related in the virilis group, including *D. montana*, *D. littoralis*, and *D. borealis* (Figure 5A & C). Outside of the virilis clade, AAACTAC is absent from the genome. We therefore searched for other satellites present in the genome using bioinformatic approaches, since no previous studies have characterized satellite DNAs in these species. Only short read genomic data were available for these species, so we characterized simple satellite DNAs with k-seek (Wei et al. 2014) and complex satellite DNAs with RepeatExplorer (Novák et al. 2013). We then selected the most abundant DNA sequence we found in each species to target with FISH (AAAC in *D. montana*, a 137bp sequence in *D. littoralis*, and ACAGACGG in *D. borealis*). First, we performed DNA FISH to verify whether the satellite sequences we characterized were indeed centromeric or pericentromeric. All three sequences were found to be in the centromeric or pericentromeric region of at least one pair of chromosomes (Figure S5). We then proceeded with RNA FISH experiments. For *D. montana*, our probe against *AAAC* again cross-hybridized with the poly-A tail and we did not detect any oocyte expression (Figure S6). However, we detected weak signal using probe against the reverse sequence *GTTT* in the late oocyte (Figure 5C). Notably, this sequence is possibly related to AAACTAC in *D. virilis*, but the expression pattern is not conserved. However, we cannot eliminate the possibility that cross-hybridization with poly-A tails may have hidden a different pattern of expression. However, both *sat-137* in *D. littoralis* and *ACAGACGG* in *D. borealis* were expressed in egg chambers. Interestingly, in these two species, transcripts were present both in nurse cell nuclei and in oocyte nuclei, differing from virilis clade species within which expression was largely isolated to the oocyte nucleus and was never found in the nurse cells.

### *D. ananassae* lncRNA candidate is expressed in nurse cells

We identified another stand-alone lncRNA in the originally-analyzed RNAseq dataset in *D. ananassae CCUGUCUGUCUGU.* We performed RNA FISH on *D. ananassae* ovaries and found that this transcript is indeed expressed, however its expression was isolated to nurse cell nuclei (Figure 5D). Furthermore, we conducted DNA-FISH to characterize the position of this satellite within the genome and found that this sequence is (peri)centromeric and present on at least one metacentric chromosome, and on multiple but not all chromosomes of *D. ananassae* (Figure S7).

## Discussion

### “Stand-alone” satellite lncRNAs are not ubiquitous in Drosophila species’ ovaries

Of the 156 satellite sequences assayed in 10 species, we only found two with an expression pattern consistent with being a stand-alone lncRNA, that is having fewer reads in the small RNA dataset compared to the total RNA dataset. We recognize that our criteria are rather stringent, since small RNA datasets are expected to be less complex, thus some of the lncRNA we found here that we classified as primarily small RNAs may still have meaningful activity as a lncRNA which can be investigated in future studies. We did indeed replicate results from (Wei et al. 2021), and found that 1.688 and Rsp were expressed at low levels as lncRNAs and high levels as piRNAs (especially 1.688). pBuM was found to be expressed in whole animals (male and female at different developmental stages) at single-digit RPM levels in *D. mojavensis* (de Lima et al. 2017). In the ovary dataset we analyzed, we found very high expression levels averaging 215 RPM and that this satellite was the second most abundantly expressed in the entire dataset (second only to AAACTAC in *D. virilis*). Small RNAs were even more abundant for this satellite, suggesting an important role of the piRNA pathway on the regulation of this satellite in *D. mojavensis* ovaries. Interestingly, past studies to date have found small RNAs for complex satellites (repeat units >20 bp), but our study showed that multiple simple satellites are also expressed as small RNAs, including AACAGAACATGTTCG in *D. simulans*, and ACAGACGG in *D. yakuba* and *D. ficusphila*. Our study demonstrates that small RNAs produced from satellite DNAs are very common, often abundant, and widespread in Drosophila, which has not been previously appreciated. These small RNAs most likely represent piRNAs, but we have not formally distinguished between piRNAs and siRNAs in our analysis. Finally, sequencing biases exist in Illumina data (Flynn et al. 2020) and it is possible that some medium to low abundance transcripts were not detected or were expressed at low, but still biologically significant levels. For example, AAACTAT was at 0.22 rpm in the total RNAseq dataset but it was still detectable with RNA FISH. The surprising result of the high expression of AAACTAC we find in *D. virilis* underscores the value of studying non-model species. Further functional studies of the *AAACUAC* transcript may lead to insights explaining why *D. virilis* expresses this lncRNA at high levels whereas other species do not express satellite lncRNAs in their oocytes.

Interestingly, we found that the two stand-alone lncRNAs were highly strand-biased with 95% or more of the reads coming from one of the two possible strands. lncRNAs that are processed into piRNAs in the *D. melanogaster* ovary have been found to be transcribed using the Rhino-Deadlock-Cutoff (RDC) complex (Wei et al. 2021), which facilitates the transcription of heterochromatic loci without canonical promoters, classically piRNA clusters. The RDC complex transcription is dual-stranded (Mohn et al. 2014), thus it is possible that the unistrand transcription of the stand-alone lncRNAs is effectuated by a different mechanism. Although the locus/loci of transcription of AAACTAC is unknown, it is unlikely to be read-through or intronic transcription from a protein-coding gene since it is expected that no genes are transcribed from the early oocyte nucleus. Non-repetitive lncRNAs are often expressed from canonical promoters (Kapusta and Feschotte 2014), but the expression mechanisms of satellite-derived stand-alone lncRNAs has not yet been investigated. However, a recent study in mouse demonstrated that inserting several tandem copies of the pericentromeric major satellite into an “inert” gene-free region of the genome was sufficient to produce transcripts, suggesting some satellites may have the “intrinsic” tendency to be expressed (Lo et al. 2026).

Upon investigating satellites in virilis group species not represented in the original RNAseq dataset, we found two other satellites (ACAGACGG in *D. borealis* and sat-137 in *D. littoralis*) that are expressed in a manner consistent with being lncRNA. We note that we did not extensively survey all possible satDNAs in *D. borealis*, *littoralis*, and *montana*; we simply tested for expression of the most abundant satellite identified from DNAseq. Although the *D. montana* sequence AAAC may share a common ancestor with AAACTAC, we found that it was mostly not expressed, except very weak expression for the reverse strand (GTTT) in late oocytes.

### *AAACUAC* has a favourable profile to investigate its potential function in oogenesis

The *AAACUAC* lncRNA is highly expressed in the *D. virilis* ovary, even higher than *bicoid*. In addition, the expression is predominantly from one strand. Finally, the expression is highly specific to the oocyte nucleus, particularly in early oogenesis when the oocyte nucleus is condensed in a specialized chromatin structure known as the karyosome and is expected to be otherwise transcriptionally silent. These findings suggest that the transcript is unlikely to be the result of generally open chromatin or permissive transcription. However, further experiments will be needed to confirm that the lncRNA is transcribed from the oocyte nucleus since there remains a possibility that it could be transcribed in nurse cells and very rapidly transported to the oocyte nucleus. We find this possibility unlikely because of the temporal expression pattern and the complete lack of signal in the nurse cells in *D. virilis* clade species. In particular, we would expect oocyte signal to increase with developmental stage if transcripts were being transported from the nurse cells, as shown for certain retrotransposons in (Wang et al. 2018). In our case, we see that expression is highest in early development, before dropping off around stage 7 and then returning around stage 9. In addition, novel mechanisms, potentially involving protein complexes, would likely be required to import lncRNAs into the nucleus of the oocyte. Regardless of the source of the transcript, we expect that future functional studies will reveal novel insights in oocyte development.

Pericentromeric satellite expression has been suggested to play a role in chromosome segregation mitosis in Drosophila cell lines (Rošić et al. 2014) and in chromatin integrity in Drosophila sperm development (Mills et al. 2019; Kumon et al. 2026). The present study is the first to show expression of a satellite lncRNA in Drosophila oogenesis. During these early stages of oogenesis, in addition to being in a condensed chromatin state, recombination has already occurred and the synaptonemal complex begins to disassemble from euchromatin while remaining intact on the pericentromeric heterochromatin (Hughes et al. 2018). It is thus possible that the *AAACUAC* lncRNA could be involved in chromosome pairing dynamics, genome stability, or chromatin maintenance. The mouse literature is relatively rich in studies about the roles of pericentromeric satellite expression (albeit at relatively lower levels than what we report here) in oogenesis and embryogenesis (Baumann et al. 2020, 2023; Probst et al. 2010). The regulation of expression and the molecular roles of the pericentromeric transcripts appears to be complex in mouse since both overexpression and knockdowns can cause defects in chromosome segregation or heterochromatin stability in different cell types (Ching et al. 2025; Baumann et al. 2023), and pericentromeric satellite transcripts can be both precursors and targets of small RNA pathways (Yadav et al. 2020; Hsieh et al. 2020; Sandoval et al. 2024). In human cells, it has recently been shown that centromeric satellite transcripts form R-loops which stabilize cohesion during cell division (Yang et al. 2026).

## Only one of two centromeric satellites are expressed

*D. virilis* chromosomes contain the pericentromeric sequence AAACTAC and centromere-proximal sequences AAACTAT (ChrY, Chr3, Chr5) and AAATTAC (ChrX, Chr2, Chr4) (Marchetti et al. 2022; Flynn et al. 2020). Interestingly, only AAACTAT is expressed and we found no detectable signal for AAATTAC. It is unclear why only one centromere-proximal satellites would be expressed, but we note two differences between these sequences based on previous studies. First, AAACTAT and AAATTAC have different structures at the DNA level, illustrated by great differences in the brightness of DAPI staining at these two sequences (Flynn et al. 2020). Despite having the exact same nucleotide composition, the order of nucleotides can impact the minor groove conformation and thus DAPI binding (Dudka et al. 2025). The different conformation of AAATTAC may prevent it from being transcribed, or it may be unstable as an RNA and rapidly degraded. Another difference between these sequences is that AAACTAT shows evidence of a recent expansion along the predicted geographic expansion of *D. virilis* (Flynn et al. 2020). An increase in the satellite’s abundance may promote expression, or the expansion could indicate that the sequence is under selection perhaps due to a (currently unknown) functional activity related to its expression. Nevertheless, expression of *AAACUAU* co-occurs with expression of *AAACUAC* in the late oocyte nucleus. Interestingly, the two transcripts mainly take up distinct regions of the nucleus. Whether the functional activity (if any) is similar between the two transcripts, or between the early and late satellite expression is unclear.

## lncRNAs outside of the virilis clade have different patterns of expression

We found that oocyte-specific expression of satellite-derived lncRNAs was restricted to virilis-clade species (*D. virilis, D. americana, D. novamexicana, D. lummei*) and the 7 nt satellite AAACTAC (and AAACTAT in *D. virilis* only), of the species and satellites we assayed. Interestingly, other virilis group species *D. littoralis* and *D. borealis* had high expression of apparently unrelated pericentromeric satellite-derived lncRNAs but the transcripts were present in both the oocyte and nurse cell nuclei. Whether these lncRNAs are transcribed from both cell types, or transcribed only in nurse cells then transported to the oocyte will have to be determined in future studies. Further, which expression pattern represents the ancestral state of this species group remains to be determined. Finally, whether the functional significance or transcriptional mechanism for the 7 nt satellites in the virilis clade versus the 8 nt and 137 nt satellites in the virilis clade remains unknown. We did not quantify any possible strand biases for these sequences. Interestingly, we found a 13 nt satellite expressed highly as a lncRNA in *D. ananassae*, but this transcript was specific to the nurse cell nuclei.

## Acknowledgements

We thank Yasir Ahmed-Braimah for discussions on this project and for sharing several fly species/stocks used in this study. We also thank Andrew G. Clark, Daniel A. Barbash, Thomas Hurd, and the Henry Krause lab for discussions and advice on data related to this project. Other Flynn lab members Laurana Germain and Philippe Dubé contributed to feedback and discussions on this project. We thank Kevin HC Wei for sharing *D. ananassae* fly stocks with us. This project was supported by startup funds from Université Laval, an NSERC Discovery grant (RGPIN-2025-04213) awarded to JMF, and a FRQNT Relève Professorale (359428) grant awarded to JMF.

## Data availability

We used already publicly-available RNAseq data. All image data is available under the accession S-BIAD3899 on the EMBL-EBI BioStudies database.

## References

1. Aditya V, Tambe V, Yue W. 2025. Development of a Novel Automated Workflow in Fiji ImageJ for Batch Analysis of Confocal Imaging Data to Quantify Protein Colocalization Using Manders Coefficient. Bio Protoc 15: e5285.

2. Adler J, Parmryd I. 2013. Colocalization analysis in fluorescence microscopy. Methods Mol Biol 931: 97–109.

3. Adler J, Parmryd I. 2010. Quantifying colocalization by correlation: the Pearson correlation coefficient is superior to the Mander’s overlap coefficient. Cytometry A 77: 733–742.

4. Bastock R, St Johnston D. 2008. Drosophila oogenesis. Curr Biol 18: R1082–7.

5. Baumann C, Ma W, Wang X, Kandasamy MK, Viveiros MM, De La Fuente R. 2020. Helicase LSH/Hells regulates kinetochore function, histone H3/Thr3 phosphorylation and centromere transcription during oocyte meiosis. Nat Commun 11: 4486.

6. Baumann C, Zhang X, Viveiros MM, De La Fuente R. 2023. Pericentric major satellite transcription is essential for meiotic chromosome stability and spindle pole organization. Open Biol 13: 230133.

7. Becalska AN, Gavis ER. 2009. Lighting up mRNA localization in Drosophila oogenesis. Development 136: 2493–2503.

8. Biscotti MA, Canapa A, Forconi M, Olmo E, Barucca M. 2015. Transcription of tandemly repetitive DNA: functional roles. Chromosome Res 23: 463–477.

9. Bolte S, Cordelières FP. 2006. A guided tour into subcellular colocalization analysis in light microscopy. J Microsc 224: 213–232.

10. Bosco G, Campbell P, Leiva-Neto JT, Markow TA. 2007. Analysis of Drosophila species genome size and satellite DNA content reveals significant differences among strains as well as between species. Genetics 177: 1277–1290.

11. Brändle F, Frühbauer B, Jagannathan M. 2022. Principles and functions of pericentromeric satellite DNA clustering into chromocenters. Semin Cell Dev Biol 128: 26–39.

12. Calvi BR, Byrnes BA, Kolpakas AJ. 2007. Conservation of epigenetic regulation, ORC binding and developmental timing of DNA replication origins in the genus Drosophila. Genetics 177: 1291–1301.

13. Ching RW, Świst-Rosowska KM, Erikson G, Koschorz B, Engist B, Jenuwein T. 2025. Forced expression of MSR repeat transcripts above a threshold limit breaks heterochromatin organisation. Nat Commun 16: 6420.

14. de Lima LG, Ruiz-Ruano FJ. 2022. In-Depth Satellitome Analyses of 37 Drosophila Species Illuminate Repetitive DNA Evolution in the Drosophila Genus. Genome Biol Evol 14. 10.1093/gbe/evac064.

15. de Lima LG, Svartman M, Kuhn GCS. 2017. Dissecting the Satellite DNA Landscape in Three Cactophilic Sequenced Genomes. G3(Bethesda) 7: 2831–2843.

16. Dudka D, Dawicki-McKenna JM, Sun X, Beeravolu K, Akera T, Lampson MA, Black BE. 2025. Satellite DNA shapes dictate pericentromere packaging in female meiosis. Nature 638: 814–822.

17. Dunn KW, Kamocka MM, McDonald JH. 2011. A practical guide to evaluating colocalization in biological microscopy. Am J Physiol Cell Physiol 300: C723–42.

18. Flynn JM, Long M, Wing RA, Clark AG. 2020. Evolutionary Dynamics of Abundant 7-bp Satellites in the Genome of Drosophila virilis. Mol Biol Evol 37: 1362–1375.

19. Flynn JM, Yamashita YM. 2024. The implications of satellite DNA instability on cellular function and evolution. Semin Cell Dev Biol 156: 152–159.

20. Gall JG, Atherton DD. 1974. Satellite DNA sequences in Drosophila virilis. J Mol Biol 85: 633– 664.

21. Gall JG, Cohen EH, Polan ML. 1971. Reptitive DNA sequences in drosophila. Chromosoma 33: 319–344.

22. Gebert D, Hay AD, Hoang JP, Gibbon AE, Henderson IR, Teixeira FK. 2025. Analysis of 30 chromosome-level Drosophila genome assemblies reveals dynamic evolution of centromeric satellite repeats. Genome Biol 26: 63.

23. Han BW, Zamore PD. 2014. piRNAs. Curr Biol 24: R730–3.

24. Hsieh C-L, Xia J, Lin H. 2020. MIWI prevents aneuploidy during meiosis by cleaving excess satellite RNA. EMBO J 39: e103614.

25. Hughes SE, Miller DE, Miller AL, Hawley RS. 2018. Female Meiosis: Synapsis, Recombination, and Segregation in. Genetics 208: 875–908.

26. Jagannathan M, Cummings R, Yamashita YM. 2018. A conserved function for pericentromeric satellite DNA. Elife 7. 10.7554/eLife.34122.

27. Kapusta A, Feschotte C. 2014. Volatile evolution of long noncoding RNA repertoires: mechanisms and biological implications. Trends Genet 30: 439–452.

28. Kinderman NB, King RC. 1973. Oogenesis indrosophila virilis. I. Interactions between the ring canal rims and the nucleus of the oocyte. Biol Bull 144: 331–354.

29. King RC, Burnett RG. 1959. Autoradiographic study of uptake of tritiated glycine, thymidine, and uridine by fruit fly ovaries. Science 129: 1674–1675.

30. Kugler J-M, Lasko P. 2009. Localization, anchoring and translational control of oskar, gurken, bicoid and nanos mRNA during Drosophila oogenesis. Fly (Austin) 3: 15–28.

31. Kumon T, Nakamizo-Dojo M, Raz AA, Lannes R, Fingerhut JM, Yamashita YM. 2026. Defective transcription of AAGAG satellite DNA causes sex-ratio meiotic drive in Drosophila. Nat Commun. 10.1038/s41467-026-72480-y.

32. Langmead B, Salzberg SL. 2012. Fast gapped-read alignment with Bowtie 2. Nat Methods 9: 357–359.

33. Li H, Handsaker B, Wysoker A, Fennell T, Ruan J, Homer N, Marth G, Abecasis G, Durbin R, 1000 Genome Project Data Processing Subgroup. 2009. The Sequence Alignment/Map format and SAMtools. Bioinformatics 25: 2078–2079.

34. Li Y, Zhang Q, Carreira-Rosario A, Maines JZ, McKearin DM, Buszczak M. 2013. Mei-p26 cooperates with Bam, Bgcn and Sxl to promote early germline development in the Drosophila ovary. PLoS One 8: e58301.

35. Lo Y-H, Shukeir N, Erikson G, Edupuganti RR, Ching R, Puri D, Jerabek L, Shiekhattar R, Jenuwein T. 2026. Transcriptional competence defines the heterochromatin nucleating potential of isolated MSR units. Nat Commun 17. 10.1038/s41467-026-70991-2.

36. Mahowald AP. 1972. Ultrastructural observations on oogenesis in Drosophila. J Morphol 137: 29–48.

37. Marchetti M, Piacentini L, Berloco MF, Casale AM, Cappucci U, Pimpinelli S, Fanti L. 2022. Cytological heterogeneity of heterochromatin among 10 sequenced Drosophila species. Genetics 222. 10.1093/genetics/iyac119.

38. Marygold SJ, Attrill H, Lasko P. 2017. The translation factors of Drosophila melanogaster. Fly (Austin) 11: 65–74.

39. Mills WK, Lee YCG, Kochendoerfer AM, Dunleavy EM, Karpen GH. 2019. RNA from a simple-tandem repeat is required for sperm maturation and male fertility in. Elife 8. 10.7554/eLife.48940.

40. Miura K, Sladoje N, eds. 2019. Bioimage data analysis workflows. 2020th ed. Springer Nature, Cham, Switzerland.

41. Mohn F, Sienski G, Handler D, Brennecke J. 2014. The rhino-deadlock-cutoff complex licenses noncanonical transcription of dual-strand piRNA clusters in Drosophila. Cell 157: 1364– 1379.

42. Navarro-Costa P, McCarthy A, Prudêncio P, Greer C, Guilgur LG, Becker JD, Secombe J, Rangan P, Martinho RG. 2016. Early programming of the oocyte epigenome temporally controls late prophase I transcription and chromatin remodelling. Nat Commun 7: 12331.

43. Ninomiya K, Yamazaki T, Hirose T. 2023. Satellite RNAs: emerging players in subnuclear architecture and gene regulation. EMBO J 42: e114331.

44. Novák P, Neumann P, Pech J, Steinhaisl J, Macas J. 2013. RepeatExplorer: a Galaxy-based web server for genome-wide characterization of eukaryotic repetitive elements from next-generation sequence reads. Bioinformatics 29: 792–793.

45. Novo CL, Wong EV, Hockings C, Poudel C, Sheekey E, Wiese M, Okkenhaug H, Boulton SJ, Basu S, Walker S, et al. 2022. Satellite repeat transcripts modulate heterochromatin condensates and safeguard chromosome stability in mouse embryonic stem cells. Nat Commun 13: 3525.

46. Palazzo AF, Koonin EV. 2020. Functional Long Non-coding RNAs Evolve from Junk Transcripts. Cell 183: 1151–1161.

47. Petrella LN, Smith-Leiker T, Cooley L. 2007. The Ovhts polyprotein is cleaved to produce fusome and ring canal proteins required for Drosophila oogenesis. Development 134: 703– 712.

48. Pezer Z, Brajković J, Feliciello I, Ugarković D. 2011. Transcription of Satellite DNAs in Insects. Prog Mol Subcell Biol 51: 161–178.

49. Poliwal B, Nikhil S, Marimuthu O, Sella RN. 2026. Uncovering the long non-coding RNAs in Drosophila: Tissue specific expressions and functional roles in reproductive development. Mol Biol Rep 53. 10.1007/s11033-026-12379-5.

50. Probst AV, Okamoto I, Casanova M, El Marjou F, Le Baccon P, Almouzni G. 2010. A strand-specific burst in transcription of pericentric satellites is required for chromocenter formation and early mouse development. Dev Cell 19: 625–638.

51. Rošić S, Köhler F, Erhardt S. 2014. Repetitive centromeric satellite RNA is essential for kinetochore formation and cell division. J Cell Biol 207: 335–349.

52. Sahoo P, Wilkins C, Yeager J. 1997. Threshold selection using Renyi’s entropy. Pattern Recognit 30: 71–84.

53. Sandoval R, Dilsavor CN, Grishanina NR, Patel V, Zamudio JR. 2024. Mammalian RNAi represses pericentromeric lncRNAs to maintain genome stability. bioRxiv. 10.1101/2024.05.09.593425.

54. Shao Z, Hu J, Jandura A, Wilk R, Jachimowicz M, Ma L, Hu C, Sundquist A, Das I, Samuel-Larbi P, et al. 2024. Spatially revealed roles for lncRNAs in Drosophila spermatogenesis, Y chromosome function and evolution. Nat Commun 15: 3806.

55. Shatskikh AS, Kotov AA, Adashev VE, Bazylev SS, Olenina LV. 2020. Functional Significance of Satellite DNAs: Insights From. Front Cell Dev Biol 8: 312.

56. van Lopik J, Alizada A, Trapotsi M-A, Hannon GJ, Bornelöv S, Czech Nicholson B. 2023. Unistrand piRNA clusters are an evolutionarily conserved mechanism to suppress endogenous retroviruses across the Drosophila genus. Nat Commun 14: 7337.

57. Wang L, Dou K, Moon S, Tan FJ, Zhang ZZ. 2018. Hijacking Oogenesis Enables Massive Propagation of LINE and Retroviral Transposons. Cell 174: 1082–1094.e12.

58. Wei KH-C, Grenier JK, Barbash DA, Clark AG. 2014. Correlated variation and population differentiation in satellite DNA abundance among lines of Drosophila melanogaster. Proc Natl Acad Sci U S A 111: 18793–18798.

59. Wei KH-C, Lower SE, Caldas IV, Sless TJS, Barbash DA, Clark AG. 2018. Variable Rates of Simple Satellite Gains across the Drosophila Phylogeny. Mol Biol Evol 35: 925–941.

60. Wei X, Eickbush DG, Speece I, Larracuente AM. 2021. Heterochromatin-dependent transcription of satellite DNAs in the female germline. Elife 10. 10.7554/eLife.62375.

61. Yadav RP, Mäkelä J-A, Hyssälä H, Cisneros-Montalvo S, Kotaja N. 2020. DICER regulates the expression of major satellite repeat transcripts and meiotic chromosome segregation during spermatogenesis. Nucleic Acids Res 48: 7135–7153.

62. Yang L, Zhang Q, Teng Z, Moran E, Lin Z, Stukenberg PT, Liu H. 2026. α-Satellite RNA maintains centromeric cohesion through R-loops. Mol Cell. 10.1016/j.molcel.2026.07.023.

63. Zafar J, Huang J, Xu X, Jin F. 2023. Recent Advances and Future Potential of Long Non-Coding RNAs in Insects. Int J Mol Sci 24. 10.3390/ijms24032605.

