## Supplementary material for "Expression of AAACTAC satellite repeats as a long noncoding RNA in the early oocyte of *Drosophila virilis*": Figure S1

**Supplementary data:**

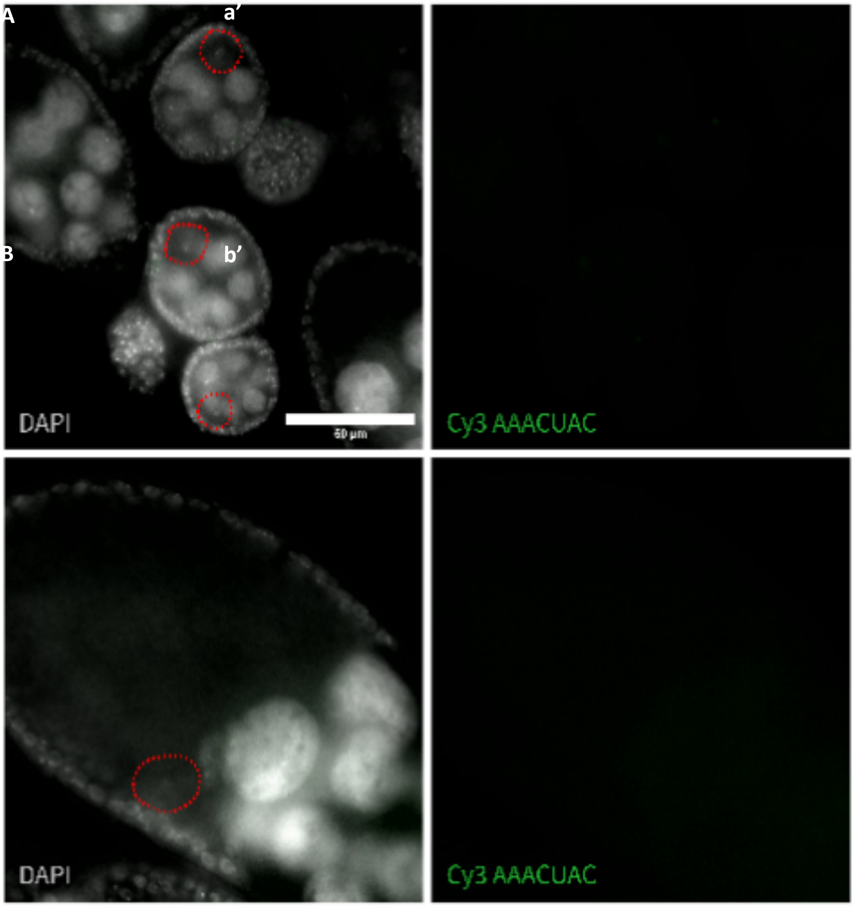


Supplementary Figure 1) A) Hybridization of the complementary probe to the forward transcript of (*AAACUAC*)n (green) after a RNAse treatment. There is a complete absence of (*AAACUAC*)n in the oocyte nucleus of stage 3-5. B) Hybridization of the complementary probe to the forward transcript of (*AAACUAC*)n (green) after a RNAse treatment, there is a complete absence of (*AAACUAC*)n in the oocyte nucleus of stage 10.

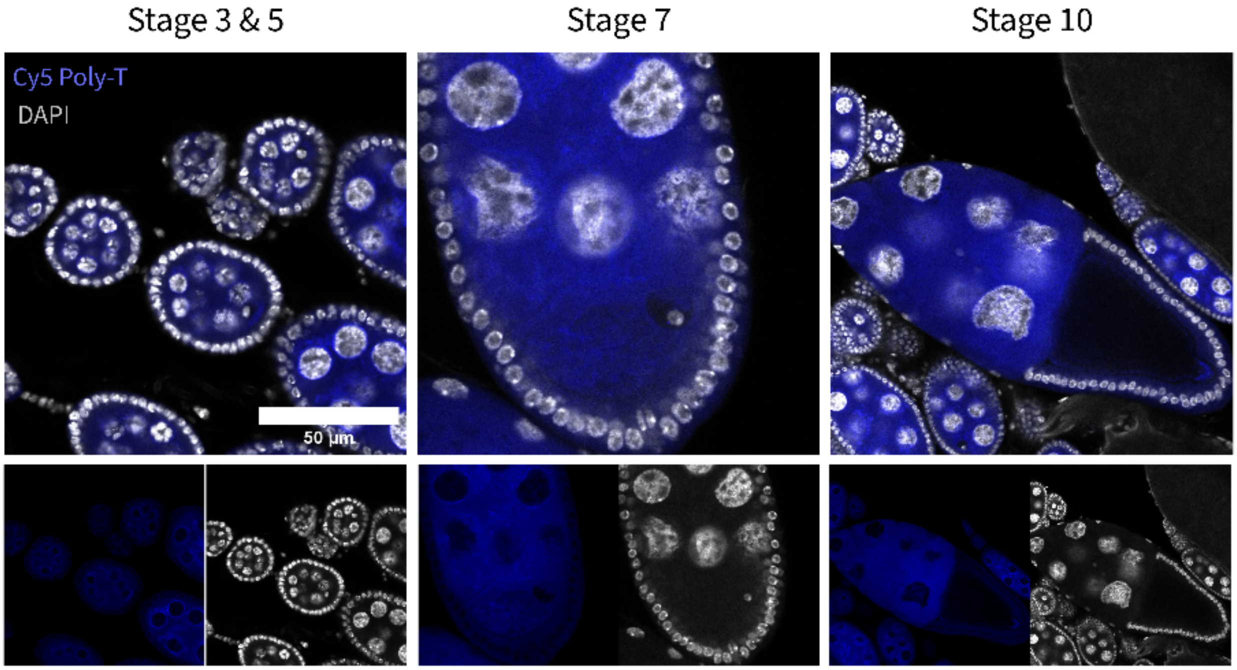


Supplementary Figure 2) RNA-Fish of the Poly-T probe in *Drosophila melanogaster*. Signal is excluded from the oocyte nucleus in early oogenesis, and some light signal in the oocyte nucleus is visible in late stages of oogenesis. Overall, the pattern matches that of *D. virilis* and is consistent with our knowledge of RNA production, transport, and localization in Drosophila oogenesis.

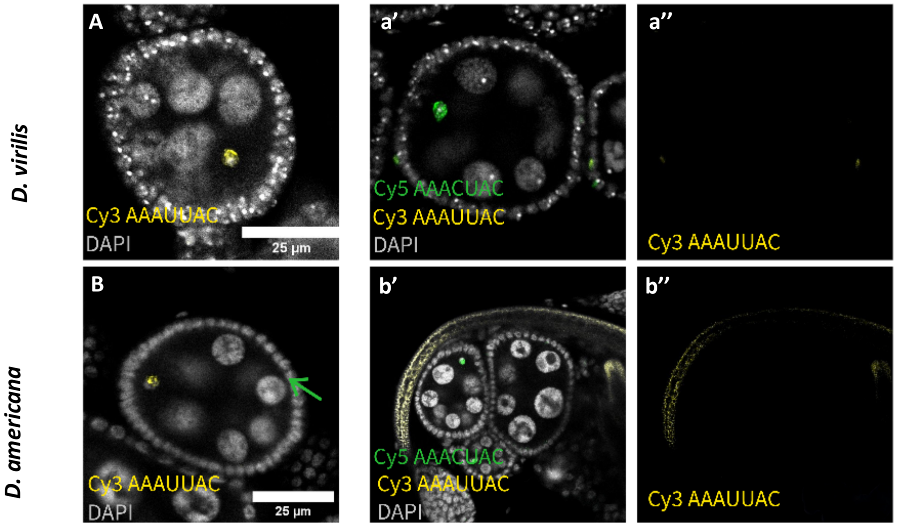


Supplementary Figure 3) A) RNA-FISH of the probe targeting *AAAUUAC* in *D. virilis* stage 5 ovarioles without multiplex A’) RNA-FISH of the probe targeting *AAAUUAC* multiplexed with the probe targeting *AAACUAC.* A’’) *AAAUUAC* channel showing no signal in the multiplex conditions. B) RNA-FISH of probe targeting *AAAUUAC* in *D. americana* stage 5 ovarioles without multiplex. *D. amerciana* does not contain the AAATTAC sequence in its genome. B’) RNA-FISH of the probe targeting *AAAUUAC* multiplexed with the probe targeting *AAACUAC* B’’) *AAAUUAC* channel showing no signal in the multiplex conditions.

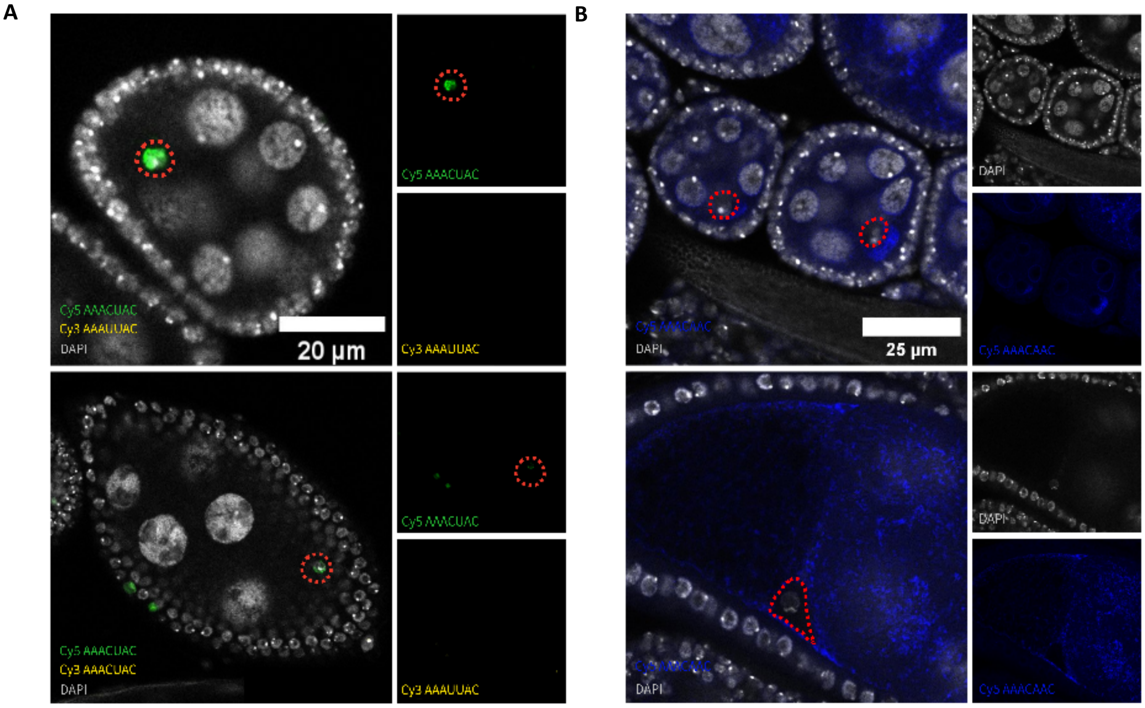


Supplementary Figure 4) A) Hybridization of the complementary probe to the probe targeting *AAAUUAC* (yellow) and *AAACUAC* (green), there is a complete absence of *AAAUUAC* in the oocyte nucleus and in the ovariole in general in *D. virilis*. B) Hybridization probe targeting AAACAAC (blue); the hybridization pattern is extremely similar to the Poly-A hybridization pattern in Figure 2. Combined with the A-rich nature of this sequence, this suggests our probe cross-hybridizes with the poly-A tail of other transcripts.

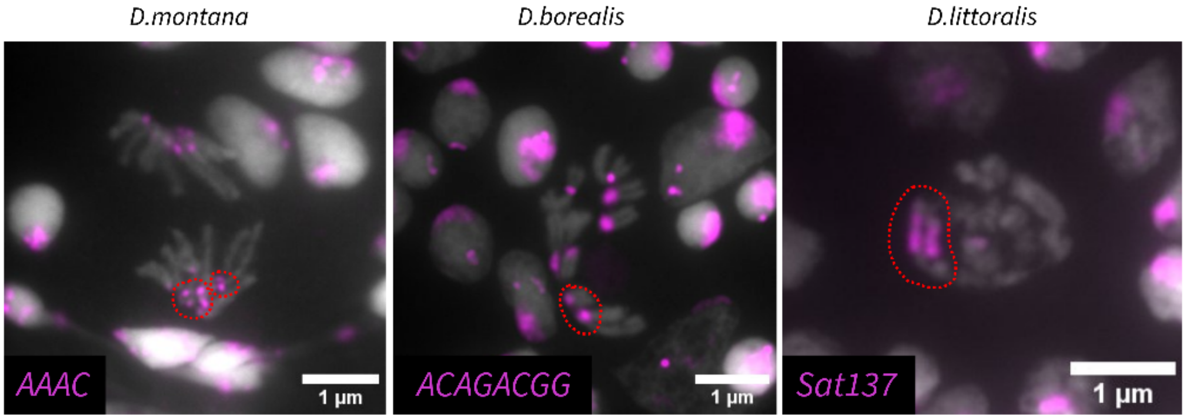


Supplementary Figure 5) DNA FISH of larval brains showing the distribution of satellite sequences specific to the three species, namely Sat137 for *D. littoralis*, (ACAGACGG)n for *D. borealis* and (AAAC)n for *D. montana*, marked in magenta. The tissues used were female larva brains.

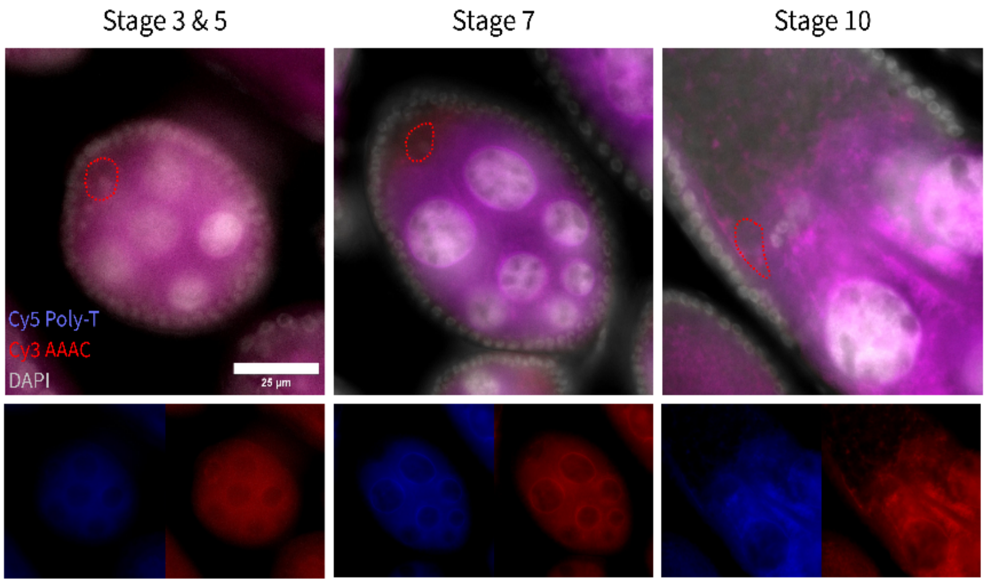


Supplementary Figure 6 shows RNA-FISH of the Poly-T probe in *Drosophila montana*, multiplexed with a probe targeting a potential AAAC transcript. We can see a strong overlap between the two signals. This indicates that AAAC likely hybridizes with the poly-A tail of mRNA.

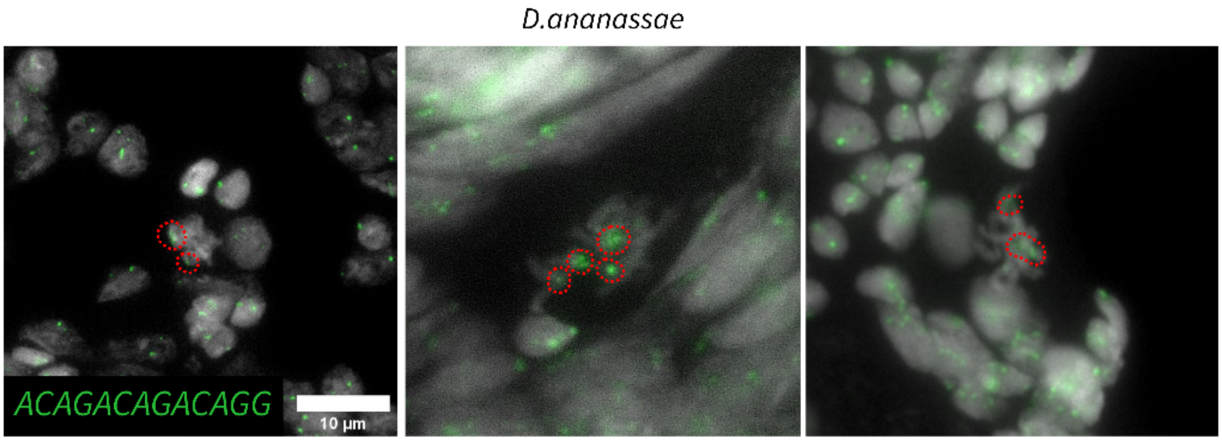


Supplementary Figure 7: DNA FISH of female larval brains showing the distribution of the satellite DNA sequence corresponding to the lncRNA we identified *D. ananassae*.

**R packages:**

| Package | Version | Citation |
|---|---|---|
| agricolae | 1.3.7 | de Mendiburu (2023) |
| car | 3.1.3 | Fox and Weisberg (2019) |
| coin | 1.4.3 | Hothorn et al. (2006); Hothorn et al. (2008) |
| DHARMa | 0.4.7 | Hartig (2024) |
| feasts | 0.5.0 | O'Hara-Wild, Hyndman, and Wang (2026) |
| gamm4 | 0.2.7 | S. Wood and Scheipl (2025) |
| ggeffects | 2.3.2 | Lüdecke (2018) |
| ggpubr | 0.6.2 | Kassambara (2025) |
| glmmTMB | 1.1.14 | Brooks et al. (2017) ; McGillycuddy et al. (2025) |
| grateful | 0.3.0 | Rodriguez-Sanchez and Jackson (2025) |
| gvlma | 1.0.0.3 | Pena and Slate (2019) |
| lmtest | 0.9.40 | Zeileis and Hothorn (2002) |
| MASS | 7.3.65 | Venables and Ripley (2002) |
| mgcv | 1.9.3 | S. N. Wood (2003) ; S. N. Wood (2004) ; S. N. Wood (2011) ; S. N. Wood, Pya, and Säfken (2016); S. N. Wood (2017) |
| multcomp | 1.4.29 | Hothorn, Bretz, and Westfall (2008) |
| nlstools | 2.1.0 | Florent Baty et al. (2015) |
| performance | 0.16.0 | Lüdecke, Ben-Shachar, et al. (2021) |
| psych | 2.6.1 | William Revelle (2026) |
| rcompanion | 2.5.0 | Mangiafico (2025) |
| see | 0.13.0 | Lüdecke, Patil, et al. (2021) |
| splines | 4.5.2 | R Core Team (2025) |
| SuppDists | 1.1.9.9 | Wheeler (2025) |
| tidyverse | 2.0.0 | Wickham et al. (2019) |
| tsibble | 1.2.0 | Wang, Cook, and Hyndman (2020) |

**3D colocalisation score model on R :**

```
model_ICQ <- lm (ICQ ~ bs (stade, 4), data = Indice_colocalisation_tout_stade)
```

**CTAC expression within all species:**

```
modele_combiné <- gam(list(score ~ sp + s (stade, by = sp, k = 10),
        ~ s (stade)),
  data = data_concaténé,
  optimizer = c("outer", "newton"),
  method = "REML",
  family = gaulss(),
  control = list(maxit = 1000))
```

**CTAC & CTAT expression in *D. virilis* :**

```
modele_combiné <— gam(list(score ~ brin + s (stade, by = brin, bs = "cr", k = 10),
              ~ s (stade)),
              data = data_combier,
              optimizer = c("outer", "newton"),
              method = "REML",
              family = gaulss(),
              control = list(maxit = 1000))
```

**Fiji :**

The version of Fiji (Schindelin et al. 2012) used is 1.54p. The plugins used during colocalization analyses are: JACoP (Bolte and Cordelières 2006), DiAna (Gilles et al. 2017) as well as CLJI2 (Haase et al. 2020). No other plugins were used to design the colocalization analysis macros/scripts.
